# Ca_V_3.1 Channels Facilitate Calcium Wave Generation and Myogenic Tone Development in Mouse Mesenteric Arteries

**DOI:** 10.1101/2023.02.10.528095

**Authors:** Mohammed A. El-Lakany, Nadia Haghbin, Naman Arora, Ahmed M. Hashad, Galina Yu. Mironova, Maria Sancho, Donald G. Welsh

## Abstract

**Background:** The myogenic response is the mechanism whereby intraluminal pressure elicits arterial constriction pursuant to the maintenance of tissue perfusion. Smooth muscle [Ca^2+^] is a key determinant of constriction, a process intimately tied to L-type (Ca_V_1.2) Ca^2+^ channels. While important, other Ca^2+^ channels, in particular T-type, are expressed and could contribute to pressure regulation within defined voltage ranges. This study examined the role of one T-type Ca^2+^ channel using mesenteric arteries from C57BL/6 wild type and Ca_V_3.1^-/-^ mice.

**Methods:** Patch-clamp electrophysiology, pressure myography, non-invasive blood pressure measurements and rapid Ca^2+^ imaging were employed to define the Ca_V_3.1^-/-^ phenotype relative to C57BL/6. Proximity ligation assay tested the closeness of Ca_V_3.1 channels to inositol triphosphate receptors (IP_3_R). Nifedipine (0.3 μM) and 2-APB (50 μM) were used to block L-type Ca^2+^ channels and IP_3_Rs, respectively.

**Results:** Initial experiments confirmed the absence of Ca_V_3.1 expression and whole-cell current in global deletion mice, a change that coincided with a reduction in systemic blood pressure. Mesenteric arteries from Ca_V_3.1^-/-^ mice produced less myogenic tone than C57BL/6, particularly at lower pressures (20-60 mmHg) where membrane potential is more hyperpolarized. This reduction in myogenic tone correlated with diminished Ca^2+^ wave generation in the Ca_V_3.1^-/-^ mice. These asynchronous events are dependent upon Ca^2+^ release from the sarcoplasmic reticulum which is insensitive to L-type Ca^2+^ channel blockade. A close physical association (<40 nm) between IP_3_R1 and Ca_V_3.1 was confirmed by proximity ligation assay; blockade of IP_3_R in nifedipine-treated C57BL/6 arteries rendered a Ca_V_3.1^-/-^ contractile phenotype.

**Conclusion:** Findings indicate that Ca^2+^ influx through Ca_V_3.1 channels contributes to myogenic tone development at hyperpolarized voltages by triggering a Ca^2+^-induced Ca^2+^ release mechanism tied to the sarcoplasmic reticulum. This study helps establish Ca_V_3.1 as a potential therapeutic target in the control of blood pressure.

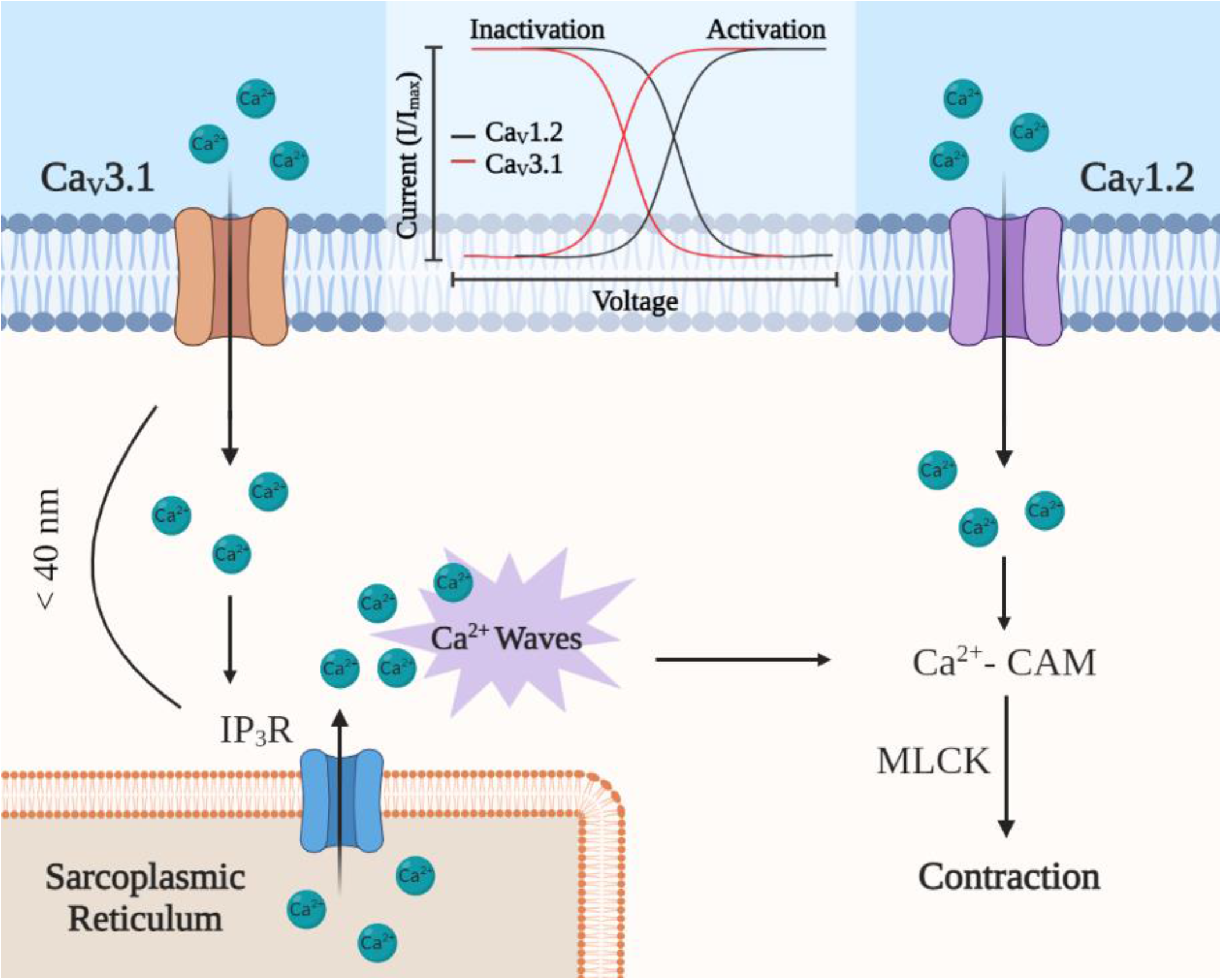

## Introduction

Smooth muscle cells in the arterial wall actively contract to intravascular pressure, maintaining organ blood flow under dynamic conditions.^1,2^ This “myogenic response” was first described by Bayliss^3^ and is intimately tied to arterial depolarization, the activation of voltage-gated Ca^2+^ channels and the concomitant rise in cytosolic Ca^2+^ concentration ([Ca^2+^]_i_), which drives myosin light chain phosphorylation.^4^ Three subclasses of voltage-gated Ca^2+^ channels (Ca_v_1-3), are expressed in the mammalian genome and each displays unique voltage-dependent properties.^5^ In arterial smooth muscle, Ca_V_1.2 (L-type) Ca^2+^ channels are principally responsible for extracellular Ca^2+^ entry and their blockade is notable for attenuating a range of constrictor responses, including those induced by pressure.^6^

L-type Ca^2+^ channels are classified as high-voltage-activated and dominate the setting of smooth muscle [Ca^2+^]_i_ when intravascular pressure is elevated and arteries depolarized.^7^ Their activity, however, markedly drops with hyperpolarization as pressure is reduced or when endothelial cell K^+^ channels are activated by selected agents.^8^ As [Ca^2+^]_i_ remains a determinant of tone, even in the hyperpolarized state, it follows that other Ca^2+^ channels, ones with a leftward voltage profile, should be expressed in vascular smooth muscle.^6^ T-type Ca^2+^ channels display activation/inactivation properties decidedly more negative to their L-type counterparts.^9,10^ Two subtypes (Ca_V_3.1 & Ca_V_3.2) are expressed in vascular smooth muscle, the latter linked to the activation of large-conductance Ca^2+^-activated K^+^ (BK) channels and a negative feedback response limiting arterial constriction.^10^ This leaves Ca_V_3.1 as to enabling myogenic tone at hyperpolarized voltages^9,11,12^, presumptively through a mechanism where Ca^2+^ influx directly contributes to the cytosolic Ca^2+^ pool or indirectly triggers sarcoplasmic reticulum Ca^2+^ release in the form of asynchronous Ca^2+^ waves.^13–15^

This study explored whether and by what mechanisms Ca_V_3.1 channels enable myogenic tone development in mouse mesenteric arteries. This work entailed the use of C57BL/6 wild type and Ca_V_3.1^-/-^ mice, and the integrated use of cellular (patch-clamp electrophysiology, immunofluorescence, and proximity ligation assay), tissue (pressure myography and rapid Ca^2+^ imaging) and whole animal (metabolic caging and blood pressure) techniques. Initial assays confirmed the absence of Ca_V_3.1 in mesenteric arterial smooth muscle of knockout animals. This absence aligned with a drop in systemic blood pressure and reduced myogenic tone at lower pressures compared to controls. Subsequent experiments revealed that Ca^2+^ wave generation was attenuated in Ca_V_3.1^-/-^ arteries as this channel no longer resided near IP_3_R1 and that IP_3_R blockade in C57BL/6 arteries produced a Ca_V_3.1^-/-^ phenotype. These findings highlight a role for Ca_V_3.1 in myogenic tone development and hemodynamic control through the triggering of sarcoplasmic reticulum Ca^2+^ waves. They additionally reveal the potential therapeutic value of Ca_V_3.1 in the control of hypertension.

## Materials and Methods

### Animal and tissue preparation

Animal procedures were approved by the animal care committee at the University of Western Ontario. Male C57BL/6 (wild type; Jackson labs) or Ca_V_3.1^-/-^ (in-house colony) mice (16-20 weeks of age) were humanely euthanized. The mesentery was removed rapidly and placed in cold PBS solution (pH 7.4) containing (in mM): 138 NaCl, 3 KCl, 10 Na_2_HPO_4_, 2 NaH_2_PO_4_, 5 glucose, 0.1 CaCl_2_, and 0.1 MgSO_4_. Third and fourth-order mesenteric were isolated and cut into 2-3mm segments and transferred to fresh cold PBS.

### Isolation of mesenteric arterial smooth muscle cells

Third and fourth-order mesenteric arteries were placed in an isolation medium (37°C, 10 minutes) containing (in mM): 60 NaCl, 80 Na-glutamate, 5 KCl, 2 MgCl_2_, 10 glucose and 10 HEPES with 1 mg/ml bovine serum albumin (BSA, pH 7.4). Vessels were then exposed to a two-step digestion process that began with 14-minute incubation (37°C) in media containing 0.5 mg/mL papain and 1.5 mg/mL dithioerythritol, followed by 10-minute incubation in media containing 100 μM Ca^2+^, and collagenases type H (0.7 mg/mL) and type F (0.4 mg/mL). Following incubation, tissues were washed repeatedly with ice-cold isolation medium and triturated with a fire-polished pipette. Liberated cells were stored in ice-cold isolation medium for use the same day.

### Immunohistochemistry

Ca_V_3.1 expression was assessed in mesenteric arterial smooth muscle cells isolated from C57BL/6 and Ca_V_3.1^-/-^ mice. Briefly, isolated cells were fixed onto a microscope cover glass in PBS (pH 7.4) containing 4% paraformaldehyde and 0.2% Tween 20. Fixed cells were blocked (1 hour, 22°C) with a quench solution (PBS supplemented with 3% donkey serum and 0.2% Tween 20) and subsequently incubated overnight (4°C, humidified chamber) with rabbit anti-Ca_V_3.1 primary antibody diluted in quench solution (1:100). In the following morning, cells were washed 3× in PBS-0.2% Tween 20 and then incubated (1 hour, 22°C) in a PBS-0.2% Tween 20 buffer containing Alexa Fluor 488 donkey anti-rabbit IgG-secondary antibody (1:200). After further washing, isolated cells and whole-mount preparations were mounted with Prolong Diamond Antifade Mountant with DAPI. Immunofluorescence was detected through a 63× oil immersion lens coupled to a Leica-TCS SP8 confocal microscope equipped with the appropriate filter sets. Smooth muscle cells isolated from C57BL/6 cerebral arteries were used as Ca_V_3.1 positive controls. Secondary antibody controls were performed and were negative for nonselective labelling.

### Electrophysiological recordings

Conventional patch-clamp electrophysiology was utilized to measure voltage-gated Ca^2+^ currents in smooth muscle cells isolated from mesenteric arteries. Cell averaged capacitance was 12–18 pF. Recording electrodes (pipette resistance, 5–8 MΩ) were fashioned from borosilicate glass using a micropipette puller (Narishige PP-830, Tokyo, Japan) and backfilled with pipette solution containing (in mM): 135 CsCl, 5 Mg-ATP, 10 HEPES, and 10 EGTA (pH 7.2). To attain a whole-cell configuration, the pipette was lowered onto a cell while applying negative pressure to rupture the membrane and garner intracellular access. Cells were voltage clamped (holding potential: −60 mV) and subjected to −90 mV followed by 10 mV voltage steps (300 ms) starting from −50 to 40 mV in a bath solution consisting of (mM): 110 NaCl, 1 CsCl, 1.2 MgCl_2_, 10 glucose, and 10 HEPES plus 10 BaCl_2_ (charge carrier). Delineation of vascular voltage-gated Ca^2+^ channels was performed by introducing 200 μM nifedipine to block L-type channels, followed by 50 μM Ni^2+^ to selectively block Ca_V_3.2 channels without affecting Ca_V_3.1. Currents were recorded using an Axopatch 200B patch-clamp amplifier (Molecular Devices, Sunnyvale, CA) at room temperature (~22°C). Data were filtered at 1 kHz, digitized at 5 kHz, and stored on a computer for offline analysis with Clampfit 10.3 software (Molecular Devices, Sunnyvale, CA). Current/voltage relationships were plotted as peak current density (pA/pF) at the different voltage steps.

### Indirect Calorimetry, Activity, and Inactivity

Comprehensive Lab Animal Monitoring System (CLAMS) interface using Oxymax software (Columbus Instruments, Columbus, OH) was utilized to measure the differences in O_2_ consumption and CO_2_ production, the cumulative amount of food and water consumed, respiratory exchange rate, activity (number of infrared beam breaks), and sleep epochs were measured in C57BL/6 and Ca_V_3.1^-/-^ mice. Chambers were kept at 24±1°C with airflow of 0.5 L/min, and animals had ad libitum access to food and water. Metabolic parameters were recorded every 10 minutes for 48 hours. Data from the same 12-hour interval for each mouse was selected to standardize the data processing.

### Blood pressure and heart rate assessment

Blood pressure measurements were performed on awake C57BL/6 and Ca_V_3.1^-/-^ mice using the non-invasive CODA tail-cuff system (Kent Scientific, CT), as described, and following recommendations of the Subcommittee of Professional and Public Education of the American Heart Association Council on High Blood Pressure Research.^16,17^ To minimize anxiety, the animals were properly acclimatized in advance, and a heating platform was used to maintain body temperature. Mice (*n* = 5) were subjected to 25-minute recordings daily for one week, and the weekly averages were recorded.

### Pressure myography

Isolated mesenteric arteries were cannulated in an arteriograph and superfused with physiological salt solution (PSS; 5% CO_2_, balance air) at 37°C containing (in mM): 119 NaCl, 4.7 KCl, 1.7 KH_2_PO_4_, 1.2 MgSO_4_, 1.6 CaCl_2_, 10 glucose, and 20 NaHCO_3_. To limit the influence of endothelial receptors, air bubbles were passed through the vessel lumen (1 min). Arterial diameters were monitored using an automated edge detection system (IonOptix, MA) and a 10× objective. Arteries were equilibrated at 15 mmHg, and contractile responsiveness was assessed by a brief (≈10 seconds) application of 60 mM KCl. After equilibration, intraluminal pressure was elevated from 20 to 100 mmHg in 20 mmHg increments for 10 minutes each, and arterial diameters were monitored in Ca^2+^ PSS (control), and in the presence of 0.3 μM nifedipine (L-type Ca^2+^ Channel blocker) alone or with 50 μM 2-APB (IP_3_R blocker). A final passive diameter assessment was conducted in Ca^2+^-free + 2 mM EGTA. Arteries that did not respond to superfused KCl (60 mM) or were insensitive or hypersensitive to pressure were excluded from experimentation. Percentage of maximum tone and incremental distensibility were calculated as follows:

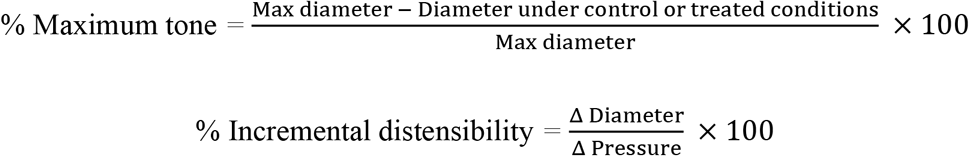

### Agonist-induced constriction

Endothelium-denuded mesenteric arteries were cannulated in a pressure myograph as explained above. Following the high potassium challenge, arteries were equilibrated at 60 mmHg, then subjected to the administration of phenylephrine (PE) into the bath solution. Increasing concentrations of PE (in M) 10^-7^, 3×10^-7^, 10^-6^, 3×10^-6^, 10^-5^, and 3×10^-5^ were superfused into the bath containing PSS in the absence (control) or presence of 0.3 μM nifedipine. Changes in diameter in response to each concentration were recorded and percentage of maximum constriction was calculated as follows:

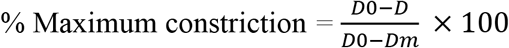

where *D* is the external diameter after each agonist concentration application, *D*0 is the external diameter in Ca^2+^ PSS, and *Dm* is the external diameter after the highest concentration of agonist under control condition.

### Calcium imaging

Freshly isolated arteries were incubated with the Ca^2+^ indicator Fluo-8 and placed on the stage of a Nikon swept-field confocal microscope with enclosed Agilent 3B laser attached to Andor camera (iXon Ultra). Fluo-8 working solution (19.1 μM) was freshly prepared by dissolving 5 μL of stock solution (1.91 mM) in 5μl pluronic acid plus 490 μL HBSS buffer consisting of (in mM): 134 NaCl, 6 KCl, 1 MgCl_2_, 2 CaCl_2_ 10 HEPES, and 10 Glucose (pH 7.4). Incubation was done for 75 minutes at 37°C in the dark. Vessels were then cannulated and equilibrated at an intraluminal pressure of 15 mmHg for 15 minutes in Ca^2+^ PSS solution. Intraluminal pressure was then raised to 60 mmHg, and Ca^2+^ waves were recorded in the presence and absence of 0.3 μM nifedipine or 50 μM 2-APB. Fluo-8-loaded arteries were excited at 488 nm and emission spectra at 510 nm viewed through a 60× water immersion objective (1.2 WI) and were monitored and analysed using Nikon NIS Elements software (AR 4.20.01). To limit laser-induced tissue injury, image acquisition was set to 45 seconds at 5 fps. A series of regions of interest (1×1 μm), created within the analysis software, was placed on 10 successive cells that were in sharp focus using the first visibly loaded smooth muscle cell as a starting point. A Ca^2+^ wave was defined as local fractional fluorescence (F/F_0_) increase above the noise level of 1.1, which spans the whole cell and lasts for at least 1 second. Ca^2+^ waves were assessed by the number of firing cells in an array of the 10 adjacent cells and the frequency of Ca^2+^ waves propagation per cell per minute.

### Proximity ligation assay

To test the spatial proximity of Ca_V_3.1 and IP_3_R, the Duolink in situ proximity ligation assay (PLA) detection kit was employed as previously described.^18^ Briefly, freshly isolated mesenteric arterial smooth muscle cells underwent successive steps of fixation (10% paraformaldehyde in PBS, 15 minutes), permeabilization (0.2% Tween 20 in PBS, 15 minutes) and blocking (Duolink blocking solution, 1 hr). Cells were then washed with PBS then incubated with primary antibodies (anti-Ca_V_3.1, anti-IP_3_R1) in Duolink antibody diluent solution at 4°C overnight. Cells were then incubated with secondary Duolink PLA PLUS and MINUS probes for 1 hour at 37°C. If target proteins are within 40 nm of each other, synthetic oligonucleotides attached to the probes hybridize enabling their subsequent amplification and binding to complementary fluorescent oligonucleotide sequences, detected using Leica-TCS SP8 confocal microscope.

### Statistical analysis

Data are expressed as means ± SEM, and *n* indicates the number of cells, arteries, or animals. Power analysis was performed *a priori* to assess the sample size sufficient for obtaining statistical significance. No more than 1 experiment was performed on cells/tissues from any given animal. Where appropriate, paired/unpaired t-tests or one-way analysis of variance (ANOVA) were performed to ascertain significant differences in mean values to a given condition/treatment. P values ≤0.05 were considered statistically significant.

### Solutions and chemicals

Fluo-8 was acquired from Abcam. Primary antibodies against Ca_V_3.1 and IP_3_R1 were purchased from NovusBio and Alomone Laboratories, respectively. Secondary antibody, Alexa Fluor 488 Donkey Anti-Rabbit IgG (H+L), and 2-APB were obtained from ThermoFisher. Duolink PLA detection kits, nifedipine, PE hydrochloride, DAPI, donkey serum and all other chemicals were obtained from Sigma-Aldrich unless stated otherwise. In cases where DMSO was used as a solvent, the maximal DMSO concentration after application did not exceed 0.5%. Please see the Major Resources Table in the Supplemental Materials

## Results

### Characterization of Ca_V_3.1^-/-^ phenotype

Genetic deletion of Ca_V_3.1 channels was confirmed by immunohistochemical analysis and the use of conventional whole-cell patch-clamp electrophysiology. In detail, Figure 1A shows Ca_V_3.1 protein expression is punctate and localized to the plasma membrane of smooth muscle cells isolated from C57BL/6 but not Ca_V_3.1^-/-^ mesenteric arteries (n=4 mice per group). This analysis aligned with whole-cell electrophysiology, which noted dampened Ca_V_3.1 activity in smooth muscle cells from Ca_V_3.1^-/-^ mice relative to C57BL/6 controls. Note, Ca_V_3.1 activity was measured by first monitoring the total inward Ba^2+^ current, the collective sum of Ca_V_1.2, Ca_V_3.1, and Ca_V_3.2 currents.^19^ Based on past studies, nifedipine and Ni^2+^ were then applied to abolish Ca_V_1.2 (L-type) and Ca_V_3.2 (T-type) activities, respectively, and the residual current was then assigned to Ca_V_3.1.^20,21^ The current-voltage relationship of each Ca^2+^ channel is illustrated in Figure 1B, C (Ca_V_3.1 current in green), with peak current (at +10 mV) summarized in Figure 1D. Recordings were attained from mesenteric smooth muscle cells (9 cells per group) isolated from 6 C57BL/6 and 8 Ca_V_3.1^-/-^ mice.

**Figure 1.**
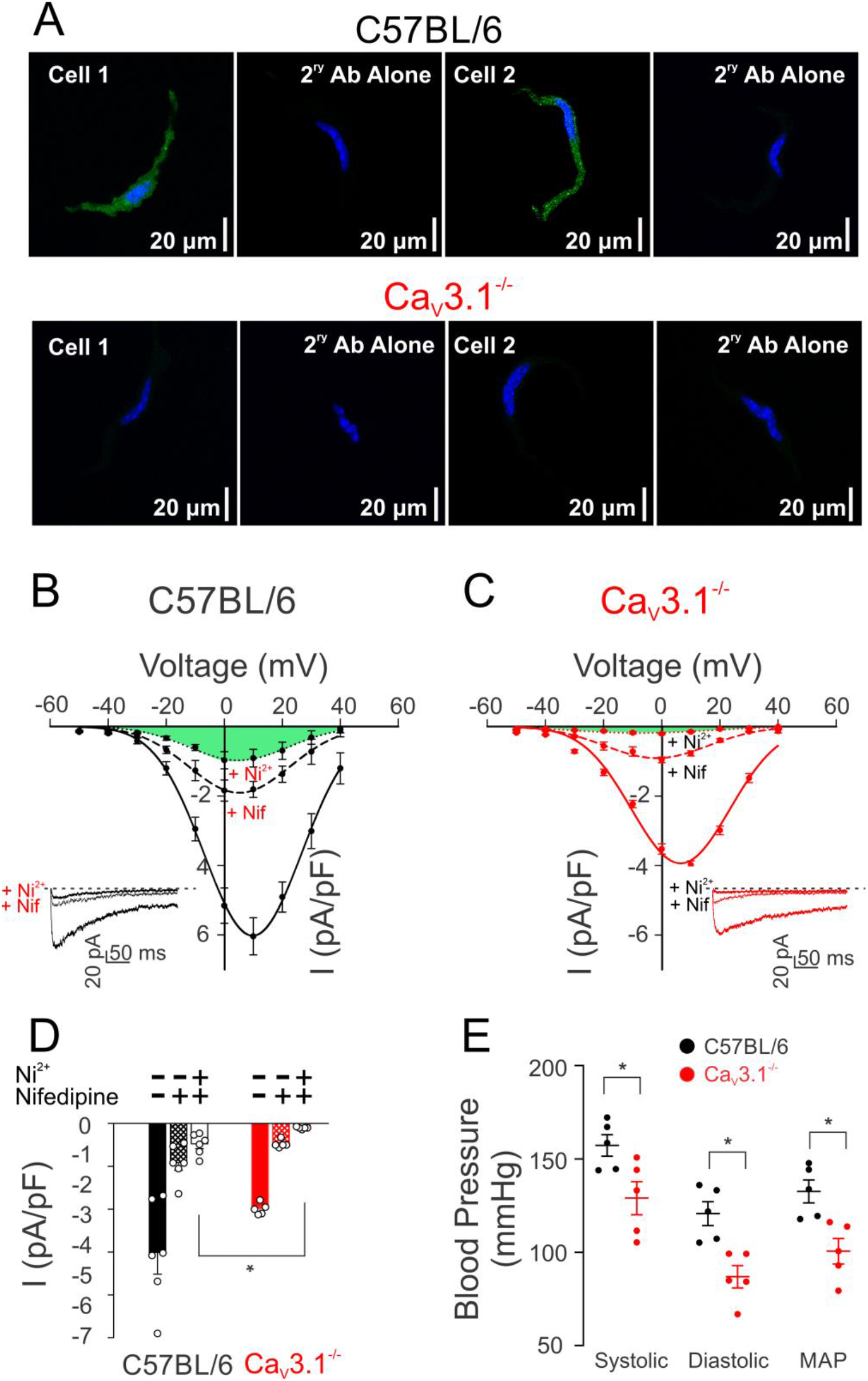
Absence of Ca_V_3.1 expression and current in SMCs isolated from mesenteric arteries of Ca_V_3.1^-/-^ mice, and lower arterial blood pressure indices in Ca_V_3.1^-/-^ mice compared to C57BL/6. **(A)** Ca_V_3.1 (green) in cerebral arterial myocytes from control mice with nuclei stained with DAPI (blue) detected with immunohistochemistry. This signal was absent in Ca_V_3.1^-/-^ mice. Secondary antibody controls were negative for nonselective labelling (*n* = 4 cells pooled from 4 animals/group). **(B)** Averaged CaV currents were assessed by whole-cell patch clamp in C57BL/6 cells showing a residual current remaining (highlighted green) after blocking L-type and Ca_V_3.2 currents by nifedipine and Ni^2+^, respectively. **(C)** Recordings of whole-cell CaV currents in Ca_V_3.1^-/-^ cells showing no residual current after nifedipine and Ni^2+^ treatment. **(D)** Peak current (I) plots of whole-cell Ba^2+^ (10 mmol/L) current before and after the application of nifedipine to C57BL/6 and Ca_V_3.1^-/-^ smooth muscle cells. (*n* = 9 SMCs from 6 mice in control group and *n* = 9 SMCs from 8 mice in knockout group. **(E)** Systolic, diastolic, and mean arterial pressure (mmHg) of Ca_V_3.1^-/-^ and C57BL/6 mice were measured using the CODA6 tail-cuff system. 25-minute recordings daily for one week were performed on both groups (*n* = 5). * indicates significance of P<0.05.

### Metabolic and blood pressure measurements in C57BL/6 and Cav3.1^-/-^ mice

Metabolic caging assessed O_2_ consumption, CO_2_ production, the respiratory exchange rate, along with cumulative food and water consumed during normal activity. No significant difference was observed among C57BL/6 and Ca_V_3.1^-/-^ mice (Table 1) in these parameters. However, Ca_V_3.1^-/-^ mice displayed a disrupted night-time sleeping pattern and were modestly but significantly heavier than the C57BL/6 controls. Subsequent tail-cuff measurements revealed that systolic, diastolic, and consequently mean arterial pressures were reduced in Ca_V_3.1^-/-^ mice compared to C57BL/6 mice (Figure 1E).

**Table 1.**
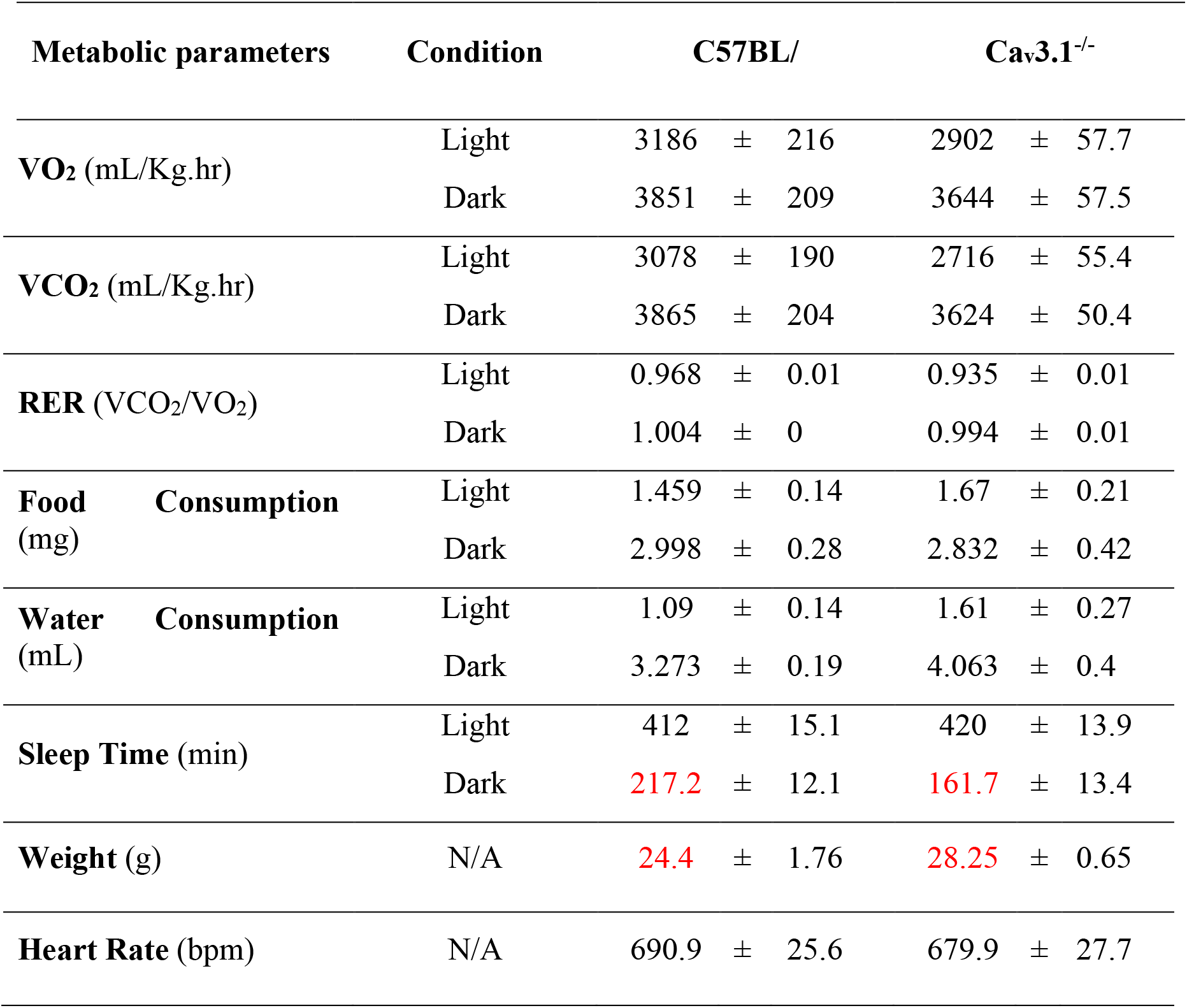
There are no discernible metabolic differences among strains, except sleep time and weight. C57BL/6 and Ca_V_3.1-/- mice were placed into individual CLAMS metabolic chambers for 48 hours. Metabolic parameters including VO_2_, VCO_2_, respiratory exchange rate (RER), food and water consumption, and sleep times, in both light and dark conditions were measured. (*n*=6 for each group). Data were presented as means ± SE and compared using unpaired two-tailed t-tests. Numbers in red indicate a significant difference of P<0.05.

### Ca_V_3.1 channels contribute to the myogenic response

Mesenteric arteries from C57BL/6 and Ca_V_3.1^-/-^ mice were mounted in a myograph and exposed to increasing intraluminal pressures (20 to 100 mmHg) in physiological saline solutions, with Ca^2+^ and Ca^2+^-free + 2 mM EGTA. Traces and summative data are presented in Figure 2A-C, and findings reveal that Ca_V_3.1^-/-^ arteries displayed reduced myogenic tone compared to C57BL/6 controls, a trend that was statistically significant at lower intraluminal pressures (20-60 mmHg). Arterial distensibility, a surrogate of vessel stiffness and defined as the percentage change in passive arteriolar diameter per change in intravascular pressure,^22^ was comparable among the two groups of arteries (Figure 2D). Control experiments using PE as a vasoconstrictor noted a comparable vasomotor tone among C57BL/6 and Ca_V_3.1^-/-^ arteries across a full concentration range (Figure 3A, B). This statement applies equally to tone generated in the absence and presence of nifedipine, except at the higher agonist concentrations where the L-type Ca^2+^ channel blocker initially appeared to be less impactful in Ca_V_3.1^-/-^ arteries (Figure 3A, B). Note, however, when this data was normalized to the % maximal constriction, the nifedipine-sensitive and insensitive components of agonist-induced constriction were comparable among the two groups of arteries (Figure 3C).

**Figure 2.**
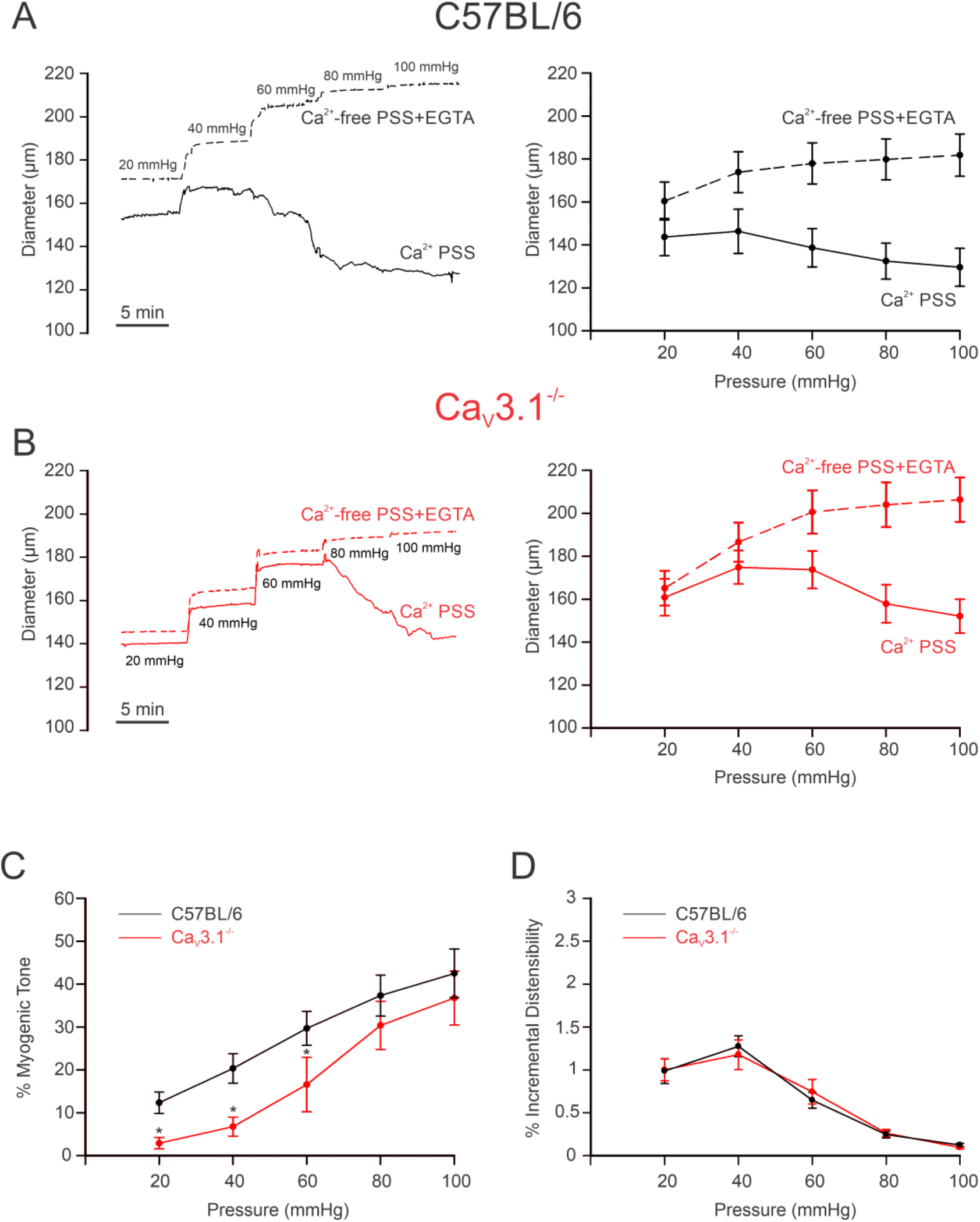
Arteries from Ca_V_3.1^-/-^ mice develop less myogenic tone at low intraluminal pressures. Isolated mesenteric arteries from Ca_V_3.1^-/-^ and C57BL/6 mice underwent a pressure curve in two conditions: PSS containing Ca^2+^ and Ca^2+^ free PSS + 2 mM EGTA, a Ca^2+^-chelating agent. A, B) Representative trace and summary data of changes in diameter in response to pressure curve (20-100 mmHg) in C57BL/6 and Ca_V_3.1^-/-^. C) Summary data shows arteries from Ca_V_3.1^-/-^ mice had lower myogenic tone in the pressure range from 20-60. D) Summary data of incremental distensibility shows no difference between groups. (*n*=6 arteries from 6 animals for each experiment). * indicates significance of P<0.05.

**Figure 3.**
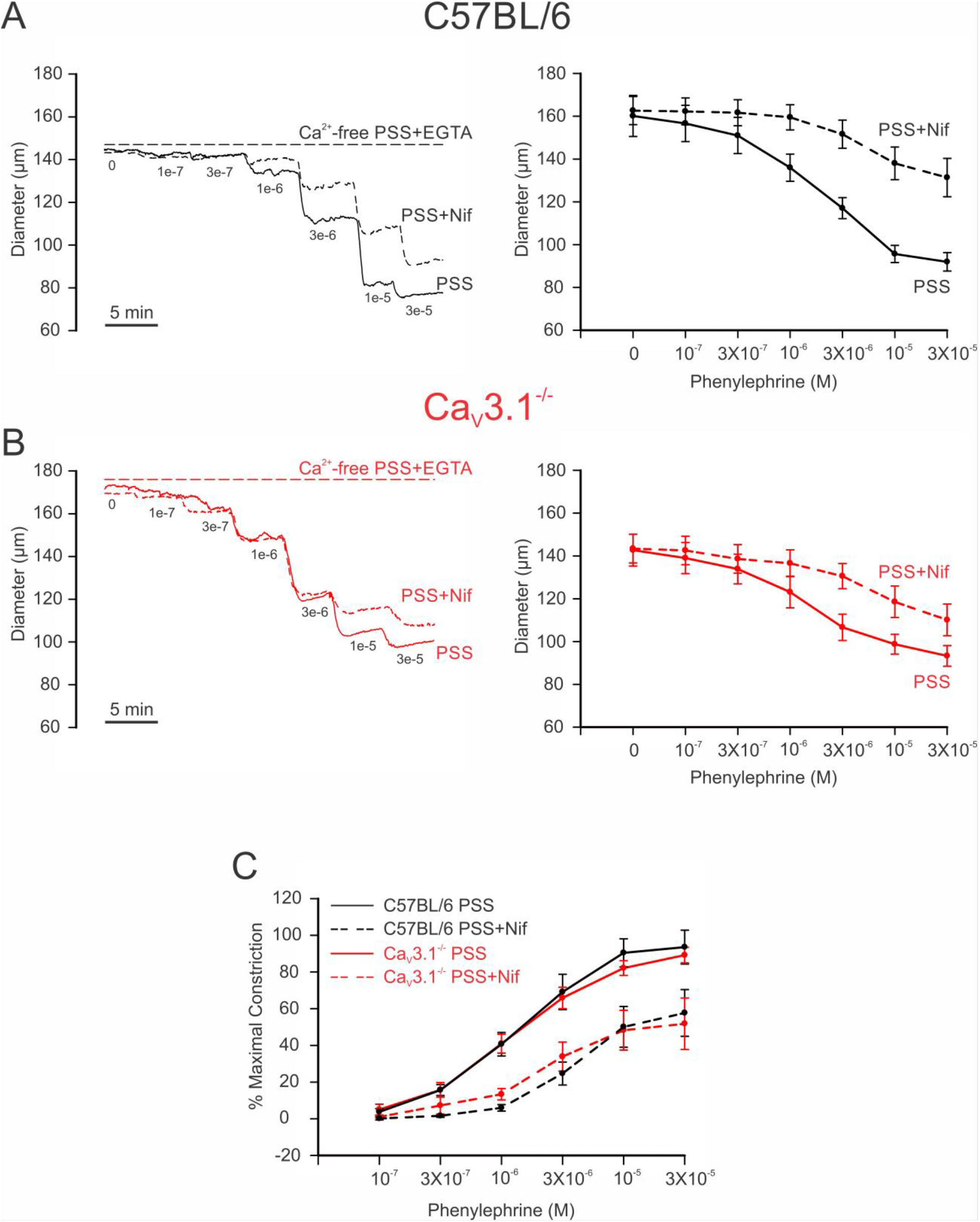
Ca_V_3.1 deletion has no impact on phenylephrine-induced constriction. Increasing concentrations of phenylephrine were applied onto pressurized arteries isolated from C57BL/6 and Ca_V_3.1^-/-^ mice in the presence and absence of nifedipine (L-type Ca^2+^ channel blocker). Experiments were conducted at an intraluminal pressure of 60 mmHg. **(A, B)** Representative traces (Left) and summary data (Right) of changes in diameter in response to phenylephrine showing a decrease in constriction in nifedipine-treated vessels from both strains. **(C)** % Maximal phenylephrine-induced constriction relative to KCl-induced constriction shows no significant difference in agonist-induced constriction between C57BL/6 and Ca_V_3.1^-/-^ mice. (*n*=6 arteries from 6 animals).

### Ca_V_3.1 enable myogenic tone by facilitating Ca^2+^ wave generation

To assess whether Ca^2+^ flux through Ca_V_3.1 triggers Ca^2+^ wave generation, mesenteric arteries from C57BL/6 and Ca_V_3.1^-/-^ mice were loaded with Fluo-8, and rapid Ca^2+^ imaging was assessed by swept field confocal microscopy. Ca^2+^ waves in C57BL/6 mice were readily observed in 80% of smooth muscle cells (8 of 10 per vessel) at a frequency of 9 waves/cell/min, each with a duration of 3-4 sec (Fig 4A, B). Similar to rat vessels, nifedipine application had little discernible effect on Ca^2+^ wave generation.^23^ The deletion of Ca_V_3.1 markedly reduced the number of firing smooth muscle cells along with firing frequency by 55% and 65%, respectively (Fig 4A,B); the Ca^2+^ waves that remained were insensitive to nifedipine. Control experiments in C57BL/6 mesenteric arteries (Figure 4C, D) subsequently confirmed that 2-APB, a blocker of IP_3_Rs, notably attenuated the number of firing cells and Ca^2+^ wave frequency by 80% and 75%, respectively. With this functional evidence indicating that Ca^2+^ flux through Ca_V_3.1 triggers IP_3_Rs and the induction of Ca^2+^ waves, the proximity ligation assay was employed to assess whether these two proteins sat closely to one another. Consistent with Ca_V_3.1 and IP_3_R1 residing within 40 nm of one another, we observed red punctate labelling in smooth muscle cells isolated from C57BL/6 by not Ca_V_3.1 mesenteric arteries (Figure 5). Controls were performed on cells treated with anti-Ca_V_3.1, anti-IP_3_R1, or secondary antibodies alone, and revealed no evidence of nonspecific binding and false product amplification.

**Figure 4.**
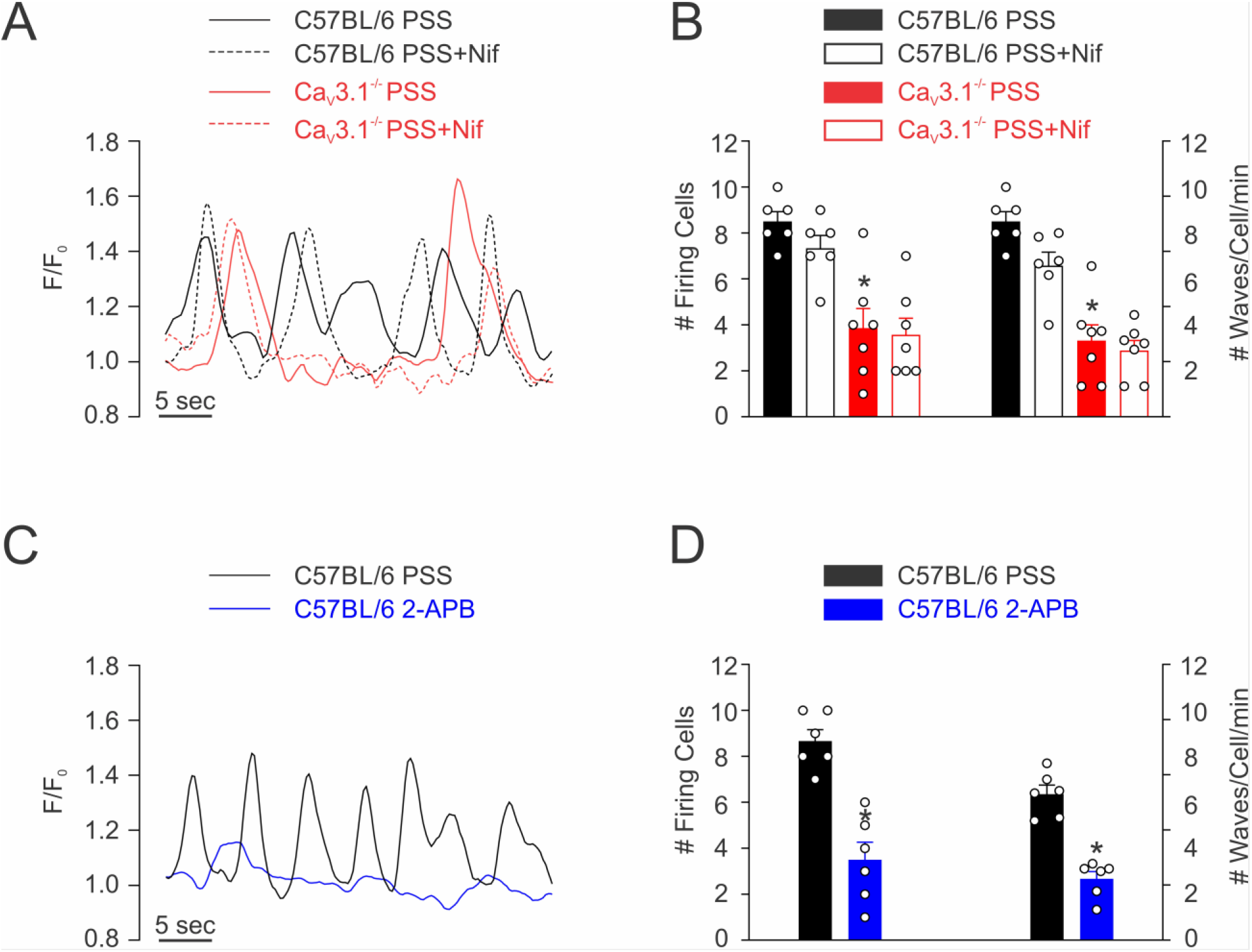
Functional roles of Ca_V_3.1 and IP_3_Rs in Ca^2+^ waves generation. Rapid Ca^2+^ imaging was performed on Fluo-8-loaded arteries from Cav3.1^-/-^ and C57BL/6 mice at an intraluminal pressure of 60 mmHg. **(A)** Representative traces from C57BL/6 and Ca_V_3.1^-/-^ mesenteric arteries with and without nifedipine. **(B)** Summary data (*n* = 6 arteries from 6 mice). Nifedipine did not impact the number of cells firing or the firing frequency in either strain. **(C)** Representative traces from C57BL/6 mesenteric arteries with and without 2-APB. **(D)** Summary data (*n* = 6 arteries from 6 mice). 2-APB (IP_3_R inhibitor) decreased the number of cells firing and their firing frequency in mesenteric arteries from C57BL/6 mice. F, fluorescence intensity; F_o_, baseline fluorescence. * indicates significance of P<0.05.

**Figure 5.**
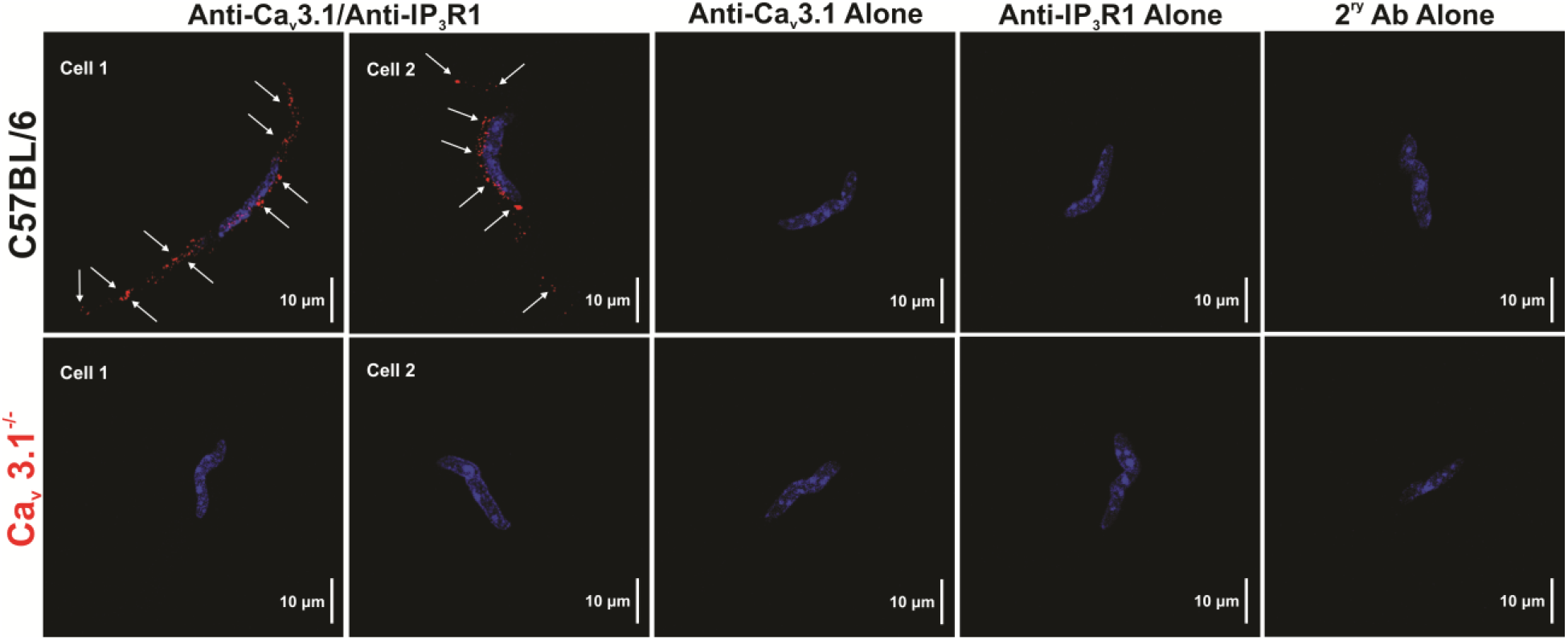
Ca_V_3.1 channels colocalize with IP_3_Rs. Proximity ligation assay was employed using isolated mesenteric arterial SMCs from C57BL/6 and Ca_V_3.1^-/-^ mice to determine the close association (<40 nm) of Ca_V_3.1 and IP_3_R1 proteins (red, denoted by white arrows). Nuclei were stained with DAPI (Blue). Control experiments used only one primary antibody or no primary antibody. (*n*=3 cells from 3 animals for each condition).

Given the preceding observation, a final set of functional experiments were performed to address the contributory role of IP_3_R-dependent Ca^2+^ waves to pressure-induced constriction. Using mesenteric arteries from C57BL/6 and Ca_V_3.1^-/-^ mice, myogenic tone was examined over a full pressure range in the absence and presence of nifedipine (0.3 μM) ± 2-APB (50 μM). Of particular note, was the nifedipine-resistant tone that was present in C57BL/6 but not Ca_V_3.1^-/-^ arteries, particularly at lower intravascular pressures (Figure 6A, B). That tone per se was largely eliminated with the further application of 2-APB, consistent with IP_3_Rs and Ca^2+^ waves playing a role in its genesis (Figure 6C). Note that IP_3_R inhibition in Ca_V_3.1^-/-^ arteries had no discernible effect on nifedipine insensitive tone at any pressure (Figure 6D).

**Figure 6.**
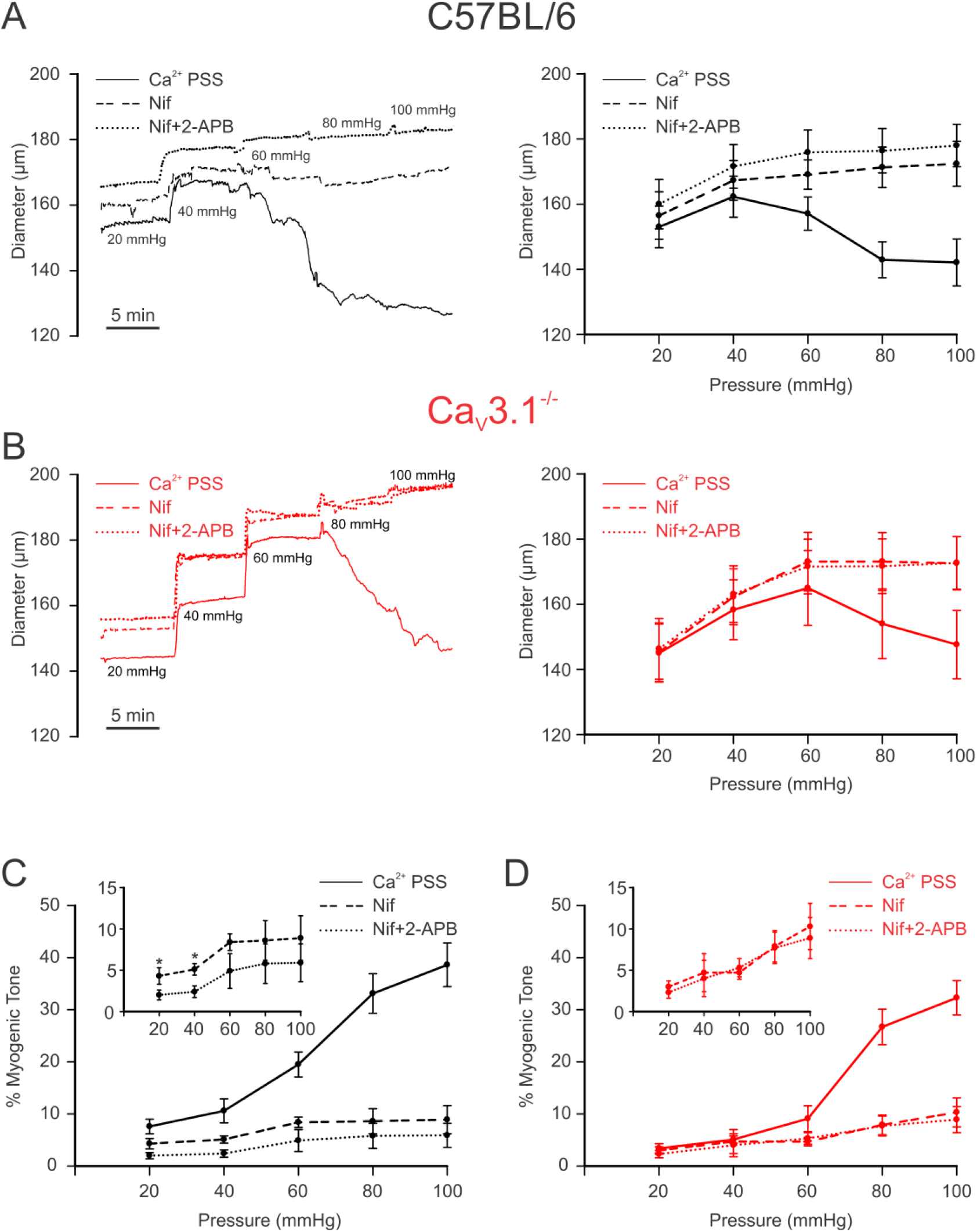
IP_3_R blockade has no impact on myogenic tone development in Cav3.1^-/-^. Mesenteric arteries isolated from C57BL/6 and Ca_V_3.1^-/-^ mice underwent stepwise pressure increases in control conditions (Ca^2+^ PSS), with nifedipine (Ca_V_1.2 blocker) alone and with 2-APB (IP_3_R blocker). **(A, B)** Representative traces (Left) and summary data (Right) of changes in mesenteric arteriolar diameter in response to pressure steps from C57BL/6 **(A)** and Ca_V_3.1^-/-^ mice **(B)** are depicted. In C57BL/6 mice, pressure-induced constriction decreased after both nifedipine and 2-APB treatment. In Ca_V_3.1^-/-^ mice, the vasomotor response following nifedipine and 2-APB treatment was not different from nifedipine treatment only. **(C)** % Myogenic tone was reduced following 2-APB treatment at 20-40 pressure range in C57BL/6 mice. **(D)** No changes in myogenic tone were observed following 2-APB treatment in Cav3.1^-/-^ mice. (*n*=6 arteries from 6 animals for each experiment). * indicates significance of P<0.05.

## Discussion

Bayliss first described the intrinsic ability of resistance arteries to constrict to a rise in intravascular pressure.^3^ This foundational response is now known to set basal tone in key organs and stabilizes organ perfusion as blood pressure fluctuates. Further, this response has been intimately tied to arterial depolarization and the rise in [Ca^2+^]_i_ enabled by graded Ca^2+^ entry principally through L-type Ca^2+^ channels.^9^ Vascular L-type Ca^2+^ channels are encoded by the Ca_V_1.2 α_1_ pore-forming subunit whose steady-state voltage-dependent properties are shifted rightward to more depolarized potentials.^24^ Recent work has revealed that L-type Ca^2+^ channels are not alone in vascular smooth muscle and that T-type Ca^2+^ channels are also expressed, with Ca_V_3.1 being key to this investigation.^9^ Its steady-state activation profile is hyperpolarized, and as such enables Ca^2+^ entry when L-type Ca^2+^ channels are deactivated. In theory, Ca^2+^ entry via T-type Ca^2+^ channels could impact tone development by directly contributing to the cytosolic Ca^2+^ pool or by acting locally and indirectly to trigger Ca^2+^ waves. Ca^2+^ waves are slow asynchronous events that spread from end to end and whose triggering is tied to the opening of sarcoplasmic reticulum IP_3_Rs by IP_3_ and Ca^2+^.^23^ Using a Ca_V_3.1^-/-^ model, we tested whether Ca^2+^ entry through this particular T-type channel does indeed facilitate myogenic tone at hyperpolarized voltages and if this functional response is coupled to the governance of Ca^2+^ waves.

### Ca_V_3.1^-/-^ model: Blood pressure and myogenic tone development

Our examination of Ca_V_3.1 began with experiments confirming the absence of Ca_V_3.1 in mesenteric arterial smooth muscle cells from genetic deletion mice (Figure 1). Two approaches were used, the first being immunohistochemistry, where surface expression of Ca_V_3.1 was notably lacking in Ca_V_3.1^-/-^ but not C57BL/6 cells. These observations aligned with whole-cell electrophysiology which revealed that the nifedipine/Ni^2+^ resistant Ba^2+^ current, previously ascribed to Ca_V_3.1^20,21^ was also absent in mesenteric arterial smooth muscle cells harvested from genetic deletion mice. The absence of this T-type Ca^2+^ channel coincided with a reduction in systolic and diastolic blood pressure, a finding consistent with a role in hemodynamic control. Past observations are limited, with one study reporting no difference in blood pressure, although values were unrealistically low for both Ca_V_3.1^-/-^ and C57BL/6 mice.^25^ A second showed blood pressure trending lower in Ca_V_3.1^-/-^ mice, along with a more substantive reduction in blood pressure variability.^26^ While the mechanism driving the blood pressure change is unclear, it’s reasonable to assert a role for diminished myogenic tone, an idea we tested in isolated mesenteric arteries across a full pressure range. Consistent with expectations, a reduction in myogenic tone was observed in Ca_V_3.1^-/-^ arteries, particularly at lower pressure (Figure 2) when vessels are hyperpolarized and T-type Ca^2+^ channels more active in the steady state.^27^ In considering these observations, prudent controls are key, the first being an assessment of an artery’s passive structural properties. In this regard, we observed no change in the arterial distensibility in vessels harvested from Ca_V_3.1^-/-^ or C57BL/6 mice. Likewise, in a second set of controls, this study did not observe a change in arterial contractility to PE across a full concentration range in the absence or presence of nifedipine, an L-type Ca^2+^ channel blocker (Figure 3). These results confirm that the molecular machinery mediating PE-induced constriction remains intact in Ca_V_3.1^-/-^ animals, as does the signalling pathways downstream from the α_1_-adrenoreceptor. Past studies have performed similar agonist controls and findings have been somewhat conflicted, with Ca_V_3.1 deletion notably reducing mesenteric arterial responsiveness in one study^28^, yet having the markedly opposite effect in another, presumptively increasing the Ca^2+^ sensitivity of the contractile apparatus.^19^

In contextualizing the preceding observations, one should recognize past inferential work linking T-type Ca^2+^ channels to myogenic tone using pharmacology with known off-target effects. This approach typically entailed the probing of myogenic tone at rest and in the presence of an L-type Ca^2+^ channel blocker to isolate residual tone whose sensitivity to T-type Ca^2+^ channel inhibition was then tested.^12,29,30^ One should also consider past work with Ca_V_3.1^-/-^ mice^31^ highlighting a role for Ca_V_3.1 channels in tone development (at low pressure), although without defining mechanism.^19^ Finally, in using this genetic deletion model, acknowledgement of other cardiovascular effects is key, in particular bradycardia^25^ and impaired blood pressure regulation through impaired NO formation.^32^

### The Role for Ca_V_3.1 in Ca^2+^ waves generation and tone development

Ca^2+^ waves are slow-spreading, end-to-end events initiated by a stimulus that drives the release of Ca^2+^ from the sarcoplasmic reticulum.^33,34^ The initiation and spread of these asynchronous events are tied to IP_3_R, Ca^2+^-permeable channels whose activation depends on IP_3_ and Ca^2+^ binding to cytosolic sites.^35^ Past work in rat cerebral arteries has shown that Ca^2+^ waves are present at low intravascular pressure and that frequency rises as pressure is elevated into the lower physiological range.^23^ Pharmacological attenuation of Ca^2+^ waves through IP_3_R blockade or impairment of store refilling results in diminished pressure-induced constriction particularly at low intravascular pressure when arteries are more hyperpolarized.^23^ Respectful of these results, it follows that low threshold Ca_V_3.1 channels provide the Ca^2+^ needed to trigger Ca^2+^ waves and foster myogenic tone when L-type Ca^2+^ channels are decidedly less active. This concept was tested three ways, the first examining Ca^2+^ wave generation in Ca_V_3.1^-/-^ arteries, the second ascertaining if Ca_V_3.1 colocalized with IP_3_Rs, and the final determining if Ca^2+^ wave inhibition in C57BL/6 mice results in a Ca_V_3.1^-/-^ contractile phenotype. Findings in Figure 4 first reveal that Ca^2+^ wave generation is robust in control mesenteric arteries as defined by the number of firing cells and the rate of Ca^2+^ waves per firing cell. Analogous to past work in rat cerebral arteries, nifedipine didn’t impact Ca^2+^ wave generation, consistent with L-type Ca^2+^ channels playing little role in initiating or maintaining these events.^23^ Ca^2+^ waves were significantly reduced in Ca_V_3.1^-/-^ arteries and abolished in control arteries by 2-APB, an IP_3_R inhibitor, findings consistent with this T-type Ca^2+^ channel driving sarcoplasmic reticulum dependent events. These intriguing findings aligned with results from the proximity ligation assay that note a close spatial association between Ca_V_3.1 and IP_3_R. In detail, this assay involves the binding of primary antibodies to two target proteins and then uses secondary antibodies with conjugated DNA strands which form a circular DNA template for amplification if proteins are <40 nm apart.^10^ The amplified product, detected as bright red puncta, is clearly visible in Figure 5, thus, it is logical to conclude that Ca^2+^ flux via Ca_V_3.1 should be sufficient to open IP_3_Rs. In light of both results, final experiments assessed whether reduced Ca^2+^ wave production in C57BL/6 vessels generate a functional phenotype akin to Ca_V_3.1^-/-^ arteries. In this regard, we monitored myogenic tone in mesenteric arteries (as a percentage; C57BL/6 and Ca_V_3.1^-/-^) at rest and following treatment with nifedipine alone or with 2-APB (Figure 6). We observed residual tone in nifedipine-treated C57BL/6 arteries but not Ca_V_3.1^-/-^ arteries, a difference that could be abolished, particularly at low intravascular pressures (20-40 mmHg) through IP_3_R blockade. These positive observations further linking Ca_V_3.1 to Ca^2+^ waves and myogenic tone align with studies showing that: 1) treatment of pressurized arteries with 2-APB decreased the myogenic response without altering global Ca^2+^ levels;^36^ and 2) 2-APB dilated hamster arteries actively producing Ca^2+^ waves.^37^

Two final points in this study require further consideration. First, while differences in myogenic tone between Ca_V_3.1^-/-^ and C57BL/6 arteries were evident at lower pressures (20-60 mmHg), the same trend was present at higher pressures, although without statistical significance. This finding is perhaps unsurprising as L-type Ca^2+^ channels rise to dominate [Ca^2+^]_i_ as arteries depolarize with pressurization. Second, while our work noted Ca^2+^ wave insensitivity to L-type Ca^2+^ channel blockade, like the cerebral vasculature^23^, it lies in contrast to cremaster arterioles where nifedipine attenuated Ca^2+^ wave formation.^38^

This discrepancy suggests there may be mechanistic uniqueness among vascular beds, which to date is unappreciated. Alternatively, one could potentially argue the higher concentration of nifedipine (1 μM) used on cremaster arteries may be blocking Ca_V_3.1 and consequently the triggering of IP_3_R.^39–41^ This perspective is consistent with electrophysiology observations noting that T-type Ca^2+^ channels are partially blocked by low micromolar nifedipine.^42,43^

## Conclusion

This study presents three key findings: First, Ca_V_3.1^-/-^ mice have lower blood pressure, and mesenteric arteries display diminished myogenic tone compared to controls. Second, immunohistochemical analysis reveals that Ca_V_3.1 lies within 40 nm of IP_3_R1, and when this arrangement is genetically disrupted, arteries generate fewer Ca^2+^ waves. Third, a pharmacological blockade of IP_3_Rs in C57BL/6 arteries produces a phenotype similar to Ca_V_3.1^-/-^ vessels, that being diminished myogenic tone at lower intravascular pressure. By establishing a clear sequential relationship between Ca_V_3.1, Ca^2+^ waves and myogenic tone, this study advances the understanding of vascular contractility and highlights a new target for therapeutic control. In this regard, one could provocatively suggest that development of selective Ca_V_3.1 blockers could be of value in the management of hypertension.

## Nonstandard Abbreviations and Acronyms

DAPI: 4’,6-diamidino-2-phenylindole
IP_3_: Inositol 1,4,5-trisphosphate
PE: Phenylephrine
PLA: proximity ligation assay
2-APB: 2-Aminoethoxydiphenyl borate

## Acknowledgments

We would like to thank all the members of our research group who contributed to this study. Graphical abstract was created with BioRender.com under an academic postdoc license. Dr DG Welsh holds the Rorabeck Chair in Molecular Neuroscience and Vascular Biology.

## Highlights

- Systemic blood pressure was lower in Ca_V_3.1^-/-^ mice, while isolated mesenteric arteries displayed diminished myogenic tone compared to C57BL/6 controls.
- Ca_V_3.1^-/-^ arteries generated fewer asynchronous Ca^2+^ waves, as the close spatial relationship between Ca_V_3.1 and IP_3_R was disrupted; IP_3_R blockade in C57BL/6 arteries produced a phenotype akin to that of Ca_V_3.1^-/-^ vessels.
- Ca_V_3.1 channels enable myogenic tone development, particularly at lower intravascular pressures, through a Ca^2+^-induced Ca^2+^ release mechanism that generates Ca^2+^ waves.

## References

1. Iadecola C, Nedergaard M. Glial regulation of the cerebral microvasculature. Nature neuroscience. 2007;10:1369–1376.

2. Longden TA, Dabertrand F, Koide M, Gonzales AL, Tykocki NR, Brayden JE, Hill-Eubanks D, Nelson MT. Capillary K+-sensing initiates retrograde hyperpolarization to increase local cerebral blood flow. Nature neuroscience. 2017;20:717–726.

3. Bayliss WM. On the local reactions of the arterial wall to changes of internal pressure. The Journal of physiology. 1902;28:220–231. doi: 10.1113/jphysiol.1902.sp000911

4. Brozovich FV, Nicholson CJ, Degen CV, Gao YZ, Aggarwal M, Morgan KG. Mechanisms of Vascular Smooth Muscle Contraction and the Basis for Pharmacologic Treatment of Smooth Muscle Disorders. Pharmacol Rev. 2016;68:476–532. doi: 10.1124/pr.115.010652

5. Snutch TP, Peloquin J, Mathews E, McRory JE. Molecular Properties of Voltage-Gated Calcium Channels. In: Voltage-Gated Calcium Channels. Boston, MA: Springer US; 2005:61–94.

6. Thorneloe KS, Nelson MT. Ion channels in smooth muscle: regulators of intracellular calcium and contractility. Canadian journal of physiology and pharmacology. 2005;83:215–242. doi: 10.1139/y05-016

7. Ghosh D, Syed AU, Prada MP, Nystoriak MA, Santana LF, Nieves-Cintrón M, Navedo MF. Calcium Channels in Vascular Smooth Muscle. Adv Pharmacol. 2017;78:49–87. doi: 10.1016/bs.apha.2016.08.002

8. Sandow SL, Senadheera S, Grayson TH, Welsh DG, Murphy TV. Calcium and Endothelium-Mediated Vasodilator Signaling. In: Islam MS, ed. Calcium Signaling. Dordrecht: Springer Netherlands; 2012:811–831.

9. Abd El-Rahman RR, Harraz OF, Brett SE, Anfinogenova Y, Mufti RE, Goldman D, Welsh DG. Identification of L-and T-type Ca2+ channels in rat cerebral arteries: role in myogenic tone development. American journal of physiology Heart and circulatory physiology. 2013;304:H58–71. doi: 10.1152/ajpheart.00476.2012

10. Harraz OF, Abd El-Rahman RR, Bigdely-Shamloo K, Wilson SM, Brett SE, Romero M, Gonzales AL, Earley S, Vigmond EJ, Nygren A, et al. Ca(V)3.2 channels and the induction of negative feedback in cerebral arteries. Circ Res. 2014;115:650–661. doi: 10.1161/circresaha.114.304056

11. Harraz OF, Visser F, Brett SE, Goldman D, Zechariah A, Hashad AM, Menon BK, Watson T, Starreveld Y, Welsh DG. CaV1.2/CaV3.x channels mediate divergent vasomotor responses in human cerebral arteries. J Gen Physiol. 2015;145:405–418. doi: 10.1085/jgp.201511361

12. Fernández JA, McGahon MK, McGeown JG, Curtis TM. CaV3.1 T-Type Ca2+ Channels Contribute to Myogenic Signaling in Rat Retinal Arterioles. Investigative Ophthalmology & Visual Science. 2015;56:5125–5132. doi: 10.1167/iovs.15-17299%JInvestigativeOphthalmology&VisualScience

13. Miriel VA, Mauban JR, Blaustein MP, Wier WG. Local and cellular Ca2+ transients in smooth muscle of pressurized rat resistance arteries during myogenic and agonist stimulation. The Journal of physiology. 1999;518 (Pt 3):815–824. doi: 10.1111/j.1469-7793.1999.0815p.x

14. Iino M, Kasai H, Yamazawa T. Visualization of neural control of intracellular Ca2+ concentration in single vascular smooth muscle cells in situ. EMBO J. 1994;13:5026–5031. doi: 10.1002/j.1460-2075.1994.tb06831.x

15. Mufti RE, Zechariah A, Sancho M, Mazumdar N, Brett SE, Welsh DG. Implications of alphavbeta3 Integrin Signaling in the Regulation of Ca2+ Waves and Myogenic Tone in Cerebral Arteries. Arteriosclerosis, thrombosis, and vascular biology. 2015;35:2571–2578. doi: 10.1161/ATVBAHA.115.305619

16. Lee VR, Barr KJ, Kelly JJ, Johnston D, Brown CFC, Robb KP, Sayedyahossein S, Huang K, Gros R, Flynn LE, et al. Pannexin 1 regulates adipose stromal cell differentiation and fat accumulation. Scientific reports. 2018;8:16166. doi: 10.1038/s41598-018-34234-9

17. Kurtz TW, Griffin KA, Bidani AK, Davisson RL, Hall JE. Recommendations for Blood Pressure Measurement in Humans and Experimental Animals. Arteriosclerosis, thrombosis, and vascular biology. 2005;25:e22–e33. doi: doi:10.1161/01.ATV.0000158419.98675.d7

18. Hashad AM, Sancho M, Brett SE, Welsh DG. Reactive Oxygen Species Mediate the Suppression of Arterial Smooth Muscle T-type Ca(2+) Channels by Angiotensin II. Scientific reports. 2018;8:3445. doi: 10.1038/s41598-018-21899-5

19. Björling K, Morita H, Olsen MF, Prodan A, Hansen PB, Lory P, Holstein-Rathlou N-H, Jensen LJ. Myogenic tone is impaired at low arterial pressure in mice deficient in the low-voltage-activated CaV3.1 T-type Ca2+ channel. Acta Physiologica. 2013;207:709–720. doi: https://doi.org/10.1111/apha.12066

20. Harraz OF, Welsh DG. Protein kinase A regulation of T-type Ca2+ channels in rat cerebral arterial smooth muscle. J Cell Sci. 2013;126:2944–2954. doi: 10.1242/jcs.128363

21. Lee J-H, Gomora JC, Cribbs LL, Perez-Reyes E. Nickel Block of Three Cloned T-Type Calcium Channels: Low Concentrations Selectively Block α1H. Biophys J. 1999;77:3034–3042. doi: https://doi.org/10.1016/S0006-3495(99)77134-1

22. Baumbach GL, Heistad DD, Siems JE. Effect of sympathetic nerves on composition and distensibility of cerebral arterioles in rats. The Journal of physiology. 1989;416:123–140. doi: 10.1113/jphysiol.1989.sp017753

23. Mufti RE, Brett SE, Tran CH, Abd El-Rahman R, Anfinogenova Y, El-Yazbi A, Cole WC, Jones PP, Chen SR, Welsh DG. Intravascular pressure augments cerebral arterial constriction by inducing voltage-insensitive Ca2+ waves. The Journal of physiology. 2010;588:3983–4005. doi: 10.1113/jphysiol.2010.193300

24. Amberg GC, Navedo MF. Calcium dynamics in vascular smooth muscle. Microcirculation. 2013;20:281–289. doi: 10.1111/micc.12046

25. Mangoni ME, Traboulsie A, Leoni AL, Couette B, Marger L, Le Quang K, Kupfer E, Cohen-Solal A, Vilar J, Shin HS, et al. Bradycardia and slowing of the atrioventricular conduction in mice lacking CaV3.1/alpha1G T-type calcium channels. Circ Res. 2006;98:1422–1430. doi: 10.1161/01.RES.0000225862.14314.49

26. Thuesen AD, Finsen SH, Rasmussen LL, Andersen DC, Jensen BL, Hansen PBL. Deficiency of T-type Ca(2+) channels Cav3.1 and Cav3.2 has no effect on angiotensin II-induced hypertension but differential effect on plasma aldosterone in mice. Am J Physiol Renal Physiol. 2019;317:F254–F263. doi: 10.1152/ajprenal.00121.2018

27. Li J, Stevens L, Klugbauer N, Wray D. Roles of molecular regions in determining differences between voltage dependence of activation of CaV3.1 and CaV1.2 calcium channels. J Biol Chem. 2004;279:26858–26867. doi: 10.1074/jbc.M313981200

28. Hansen PB, Poulsen CB, Walter S, Marcussen N, Cribbs LL, Skott O, Jensen BL. Functional importance of L-and P/Q-type voltage-gated calcium channels in human renal vasculature. Hypertension. 2011;58:464–470. doi: 10.1161/HYPERTENSIONAHA.111.170845

29. VanBavel E, Sorop O, Andreasen D, Pfaffendorf M, Jensen BL. Role of T-type calcium channels in myogenic tone of skeletal muscle resistance arteries. American journal of physiology Heart and circulatory physiology. 2002;283:H2239–2243. doi: 10.1152/ajpheart.00531.2002

30. Kuo IY, Ellis A, Seymour VA, Sandow SL, Hill CE. Dihydropyridine-insensitive calcium currents contribute to function of small cerebral arteries. J Cereb Blood Flow Metab. 2010;30:1226–1239. doi: 10.1038/jcbfm.2010.11

31. Kim D, Song I, Keum S, Lee T, Jeong M-J, Kim S-S, McEnery MW, Shin H-SJN. Lack of the burst firing of thalamocortical relay neurons and resistance to absence seizures in mice lacking α1G T-type Ca2+ channels. 2001;31:35–45.

32. Svenningsen P, Andersen K, Thuesen AD, Shin HS, Vanhoutte PM, Skott O, Jensen BL, Hill C, Hansen PB. T-type Ca(2+) channels facilitate NO-formation, vasodilatation and NO-mediated modulation of blood pressure. Pflugers Arch. 2014;466:2205–2214. doi: 10.1007/s00424-014-1492-4

33. Jaffe LF. The path of calcium in cytosolic calcium oscillations: a unifying hypothesis. Proc Natl Acad Sci U S A. 1991;88:9883–9887. doi: 10.1073/pnas.88.21.9883

34. Jaffe LF. Calcium waves. Philosophical transactions of the Royal Society of London Series B, Biological sciences. 2008;363:1311–1316. doi: 10.1098/rstb.2007.2249

35. Amberg GC, Navedo MF. Calcium dynamics in vascular smooth muscle. Microcirculation. 2013;20:281–289.

36. Potocnik SJ, Hill MA. Pharmacological evidence for capacitative Ca(2+) entry in cannulated and pressurized skeletal muscle arterioles. Br J Pharmacol. 2001;134:247–256. doi: 10.1038/sj.bjp.0704270

37. Jackson WF, Boerman EM. Regional heterogeneity in the mechanisms of myogenic tone in hamster arterioles. American journal of physiology Heart and circulatory physiology. 2017;313:H667–H675. doi: 10.1152/ajpheart.00183.2017

38. Jackson WF, Boerman EM. Voltage-gated Ca(2+) channel activity modulates smooth muscle cell calcium waves in hamster cremaster arterioles. American journal of physiology Heart and circulatory physiology. 2018;315:H871–H878. doi: 10.1152/ajpheart.00292.2018

39. McDonald TF, Pelzer S, Trautwein W, Pelzer DJ. Regulation and modulation of calcium channels in cardiac, skeletal, and smooth muscle cells. Physiological reviews. 1994;74:365–507. doi: 10.1152/physrev.1994.74.2.365

40. Curtis TM, Scholfield CN. Nifedipine blocks Ca2+ store refilling through a pathway not involving L-type Ca2+ channels in rabbit arteriolar smooth muscle. The Journal of physiology. 2001;532:609–623. doi: 10.1111/j.1469-7793.2001.0609e.x

41. Akaike N, Kanaide H, Kuga T, Nakamura M, Sadoshima J, Tomoike H. Low-voltage-activated calcium current in rat aorta smooth muscle cells in primary culture. The Journal of physiology. 1989;416:141–160. doi: 10.1113/jphysiol.1989.sp017754

42. Akaike N, Kostyuk PG, Osipchuk YV. Dihydropyridine-sensitive low-threshold calcium channels in isolated rat hypothalamic neurones. The Journal of physiology. 1989;412:181–195. doi: 10.1113/jphysiol.1989.sp017610

43. Shcheglovitov A, Zhelay T, Vitko Y, Osipenko V, Perez-Reyes E, Kostyuk P, Shuba Y. Contrasting the effects of nifedipine on subtypes of endogenous and recombinant T-type Ca2+ channels. Biochem Pharmacol. 2005;69:841–854. doi: 10.1016/j.bcp.2004.11.024

